# Resolving the 3D rotational and translational dynamics of single molecules using radially and azimuthally polarized fluorescence

**DOI:** 10.1101/2021.10.19.465033

**Authors:** Oumeng Zhang, Weiyan Zhou, Jin Lu, Tingting Wu, Matthew D. Lew

**Affiliations:** Department of Electrical and Systems Engineering, Washington University in St. Louis, Missouri 63130, USA; Center for the Science and Engineering of Living Systems, Washington University in St. Louis, Missouri 63130, USA; Institute of Materials Science and Engineering, Washington University in St. Louis, Missouri 63130, USA

**Keywords:** Single-molecule tracking, phospholipid bilayer, cholesterol condensing, rotational diffusion, lateral diffusion

## Abstract

We report a radially and azimuthally polarized (raPol) microscope for high detection and estimation performance in single-molecule orientation-localization microscopy (SMOLM). With 5000 photons detected from Nile red (NR) transiently bound within supported lipid bilayers (SLBs), raPol SMOLM achieves 2.9 nm localization precision, 1.5° orientation precision, and 0.17 sr precision in estimating rotational wobble. Within DPPC SLBs, SMOLM imaging reveals the existence of randomly oriented binding pockets that prevent NR from freely exploring all orientations. Treating the SLBs with cholesterol-loaded methyl-β-cyclodextrin (MβCD-chol) causes NR’s orientational diffusion to be dramatically reduced, but curiously, NR’s median lateral displacements drastically increase from 20.8 nm to 75.5 nm (200 ms time lag). These jump diffusion events overwhelmingly originate from cholesterol-rich nanodomains within the SLB. These detailed measurements of single-molecule rotational and translational dynamics are made possible by raPol’s high measurement precision and are not detectable in standard SMLM.

---

Beyond improving the localization accuracy ^1–6^ of singlemolecule localization microscopy (SMLM), ^7–10^ simultaneously imaging molecular positions and orientations in singlemolecule orientation localization microscopy (SMOLM) provides unparalleled insights into biochemical processes. Recent developments in SMOLM allow scientists to resolve the organization of amyloid aggregates, ^11–13^ conformations of DNA strands, ^6,14–18^ and the structure of actin networks. ^19–21^ Using point spread function (PSF) engineering, microscopists have developed various techniques ^4,6,20,22,23^ to improve measurement precision, approaching the fundamental performance limits for measuring both orientation ^24–26^ and localization. ^27,28^

However, many SMOLM techniques exhibit poor measurement precision for molecules that are oriented out of the coverslip plane (e.g., the *x*- and *y*-polarized (*xy*Pol) standard PSF ^12,29^). Others improve measurement precision by expanding the imaging system PSF significantly and thus suffer degraded localization precision and detection sensitivity for dim emitters (e.g., the Tri-spot PSF ^22^). Despite the recent developments of CHIDO ^20^ and the vortex PSF ^6^ for imaging the 3D positions and orientations of single molecules (SMs), no existing techniques exhibit sufficiently high localization precision, orientation precision, and sensitivity to SM wobble for simultaneous 3D orientation, wobble, and position tracking of SMs within lipid membranes. Inspired by symmetries within the dipole radiation pattern, we show the first experimental demonstration of radially and azimuthally polarized (raPol) fluorescence, ^24,26,30^ a variant of the y-Phi PSF, ^5^ for highly sensitive position and orientation tracking of SMs. It is implemented using a commercially available vortex (half) wave plate (VWP) and polarizing beamsplitter (PBS) within a widefield epifluorescence microscope. The raPol microscope exhibits excellent orientation estimation performance while maintaining high detection rates and localization performance comparable to those of standard polarized microscopes. We utilize the raPol PSF to study the dynamics of Nile red (NR) molecules transiently bound to supported lipid bilayers (SLBs). ^31^ Imaging NR dynamics within DPPC SLBs reveals the existence of randomly oriented binding pockets that prevent NR from freely rotating. In addition, raPol is capable of tracking simultaneously the position and orientation of NR as it explores SLBs modified by methyl-β-cyclodextrin loaded with cholesterol (MβCD-chol). As cholesterol (chol) is deposited, NR’s orientation tilts mostly perpendicular to the SLB, and its rotational diffusion is greatly reduced, but its translational diffusion dramatically increases almost four-fold. These data suggest that NR “jumps” between cholesterol-rich nanodomains within the SLB. To our knowledge, these experiments are the first measurements of how MβCD-chol affects the nanoscale chemical environments within lipid membranes at the single-molecule level.

We begin by describing a rotationally diffusing molecule using its average orientation ***μ*** = [*μ*_*x*_, *μ*_*y*_, *μ*_*z*_]^†^ = [sin *θ* cos *ϕ*, sin *θ* sin *ϕ*, cos *θ*]^†^ within a hard-edged cone of solid angle Ω (Figure 1a), where *μ*_*z*_ is parallel to the optical axis, Ω = 0 represents a rotationally-fixed molecule, and Ω = 2*π* represents a freely-wobbling molecule. Here, we assume that the SM wobbles uniformly within the cone for simplicity, and our analysis may be easily adapted for other rotational potential wells or geometries. ^32–34^

**Figure 1:**
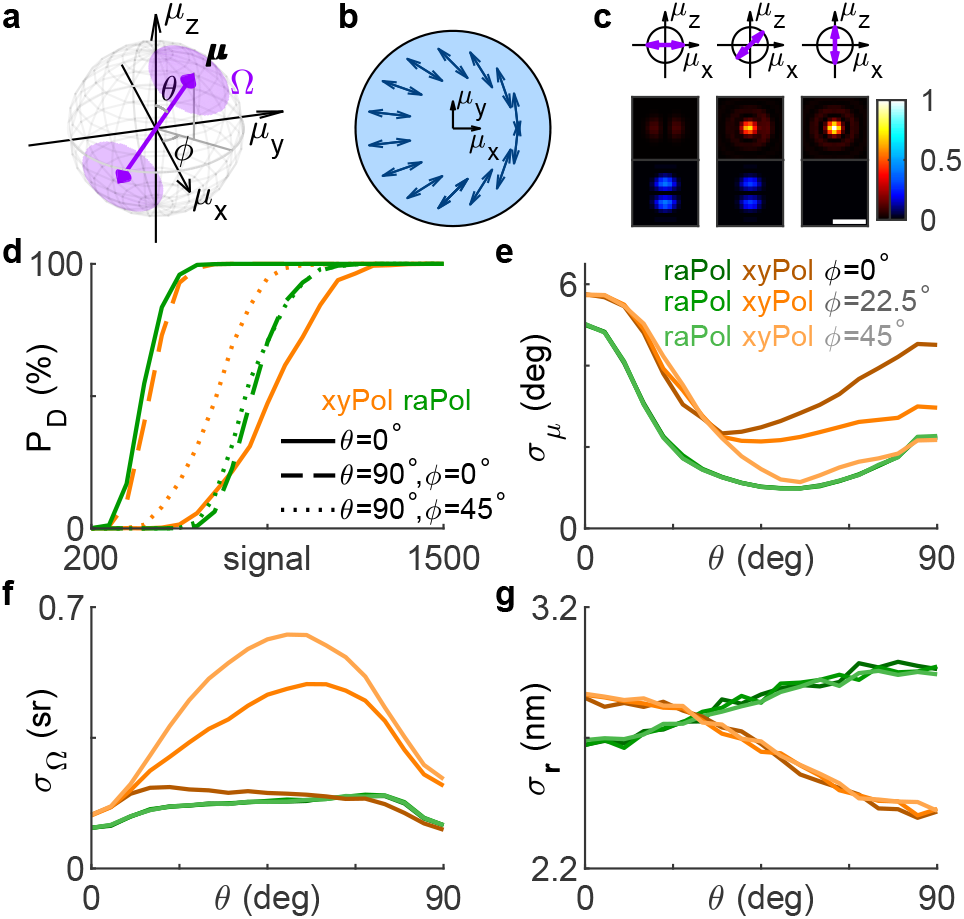
The radially and azimuthally polarized (raPol) standard PSF. (a) The rotational diffusion of a fluorescent molecule is described using a unit vector ***μ*** and a wobble solid angle Ω on the surface of a hard-edged cone. (b) A vortex (half) waveplate (VWP) is placed at the back focal plane of the imaging system, transforming radially and azimuthally polarized light to *x* and *y*-polarized light, respectively. Arrows represent the VWP’s fast axis. (c) Representative radially (red) and azimuthally (blue) polarized PSFs of rotationally-fixed molecules with polar angles *θ* of 90°, 45°, and 0°. Colorbar: normalized intensity. Scale bar: 500 nm. (d-g) Detection and estimation performance of a sparsity-promoting maximum-likelihood estimation algorithm using simulated *xy*Pol (orange) and raPol (green) images. (d) Detection rate *P*_D_ as a function of the number of total signal photons detected from an SM with various orientations and a fixed wobble Ω = 0.28*π* sr. Dashed, solid, and dotted lines represent molecular orientations *θ* = 0°, [*θ, ϕ*] = [90°, 0°], and [*θ, ϕ*] = [90°, 45°], respectively. (eg) Precision of estimating the (e) average orientation ***μ***, (f) wobble angle Ω, and (g) 2D position ***r*** of an SM with 5000 signal photons detected for various polar angles *θ*. Dark, medium, and light colors represent azimuthal angles *ϕ* = 0°, *ϕ* = 40°, and *ϕ* = 90°, respectively. The background is 30 photons per camera pixel (66.86 × 66.86 nm^2^).

Due to the symmetry of the dipole emission pattern, the photons emitted by an SM oriented parallel to the optical axis (*μ*_*z*_ = 1) captured by an objective lens are radially polarized. ^3,35,36^ Therefore, an intuitive way to distinguish out-of-plane molecules from in-plane ones is to measure radially vs. azimuthally polarized fluorescence. We achieve this polarization separation by adding a VWP (Figure 1b) to the back focal plane (BFP) of a microscope with two polarized imaging channels. The spatially-varying fast axis direction of the VWP turns the radially and azimuthally polarized light to *x* and *y*-polarized light. A PBS is then used to separate the fluorescence into radially and azimuthally polarized images (Figure S1a). As a molecule rotates out of plane, i.e., as *μ*_*z*_ increases, photons shift from the azimuthally polarized to the radially polarized channel. When the SM is aligned with the optical axis, all fluorescence photons are confined to the radially polarized image (Figure 1c, red).

High detection rates are critical for collecting unbiased measurements of SM dynamics. We quantify the detection rate *P*_D_ of raPol compared to *xy*Pol, a common easy-to-implement SMOLM technique, using RoSE-O, a sparsity-promoting joint detection and maximum-likelihood estima-tion algorithm. ^18^ We find that despite the orientation dependence, the detection performance of raPol is comparable to that of *xy*Pol (Figure 1d and Figure S3). Thus, dim SMs are detected more reliably using raPol as opposed to most engineered PSFs optimized for SMOLM; it is more difficult to detect weak emitters using larger PSFs [Figure S3c(iii,iv)]. Due to the toroidal, polarized emission pattern of an SM, the detection rate using raPol is much better than that of *xy*Pol for *z*-oriented molecules since the energy is concentrated in the radially polarized channel; with 500 signal photons detected, raPol achieves a detection rate of 92.5% while *xy*Pol can only detect less than 1% of the SMs (Figure 1d). The superior detection rate for this orientation motivates us to adopt raPol for lipid membrane imaging (see below). However, its detection rate is worse than that of *xy*Pol for inplane molecules for the opposite reason (Figure 1c). Interestingly, due to the symmetry of raPol, its detection performance is not affected by the azimuthal angle *ϕ*, whereas that of *xy*Pol may vary up to a factor of 6 across various *ϕ* (Figure S3a,b).

Next, we compare the orientation and position measurement precision for raPol and *xy*Pol, evaluated at an SBR such that we achieve 100% detection rate using both PSFs. The precision of measuring the average orientation *σ*_***μ***_ is quantified as the angle subtended by an arc that represents measurement uncertainty on the orientation unit sphere (SI Section 2.2). Strikingly, raPol exhibits superior orientation and wobble measurement precision over *xy*Pol for almost all possible SM orientations [Figure 1e,f and Figure S5a,b(ii,v)]. The average 2D localization precision *σ*_***r***_ using raPol is only 12% worse compared to that of *xy*Pol [Figure 1g and Figure S5a,b(i)], where *xy*Pol is usually perceived to be optimal for localizing in-focus molecules. Further Cramér-Rao bound (CRB, SI Section 2.2) ^37^ analysis shows that the orientation performance of raPol is comparable to recently developed engineered PSFs while exhibiting a much better localization precision (Figure S4 and Table S1).

While the translational dynamics of fluorophore-lipid interactions have been characterized at the SM level, e.g., by fluorescence correlation spectroscopy (FCS), ^41^ relatively little is known about their rotational dynamics. We next study the rotational dynamics of Nile red (Figure 2a) within SLBs ^31^ (Figure 2b) using points accumulation for imaging in nanoscale topography (PAINT). ^10^ We form a DPPC [di(16:0)PC (phosphatidylcholine), Figure S8] bilayer on coverglass and use RoSE-O to analyze images of blinking NR molecules and extract their positions and orientations. First, we measure NR within a DPPC SLB using both *xy*Pol and raPol, excited by elliptically polarized epi (Figure 2c) and single polarization total internal reflection fluorescence (TIRF, Figure 2f) illumination. Interestingly, NR’s polar orientation *θ* changes significantly with illumination polarization, from a median of 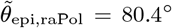 to 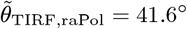, implying that NR’s emission dipole orientation is correlated with the orientation of its absorption dipole moment within a DPPC SLB. To confirm, we changed the laser beam tilt to *α* = 25° and *α* = 45° relative to the optical axis (Figure 2d,e). As the *z*-polarized electric field in the excitation beam increased, the observed NR polar angles systematically decreased 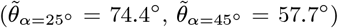. The change in polar angle is obvious within the raPol images at each illumination angle *α*; fluorescence photons become increasingly radially polarized as *α* increases [Figure 2(i)]. These trends were consistently observed across ∼35k localizations and 12 fields of view.

**Figure 2:**
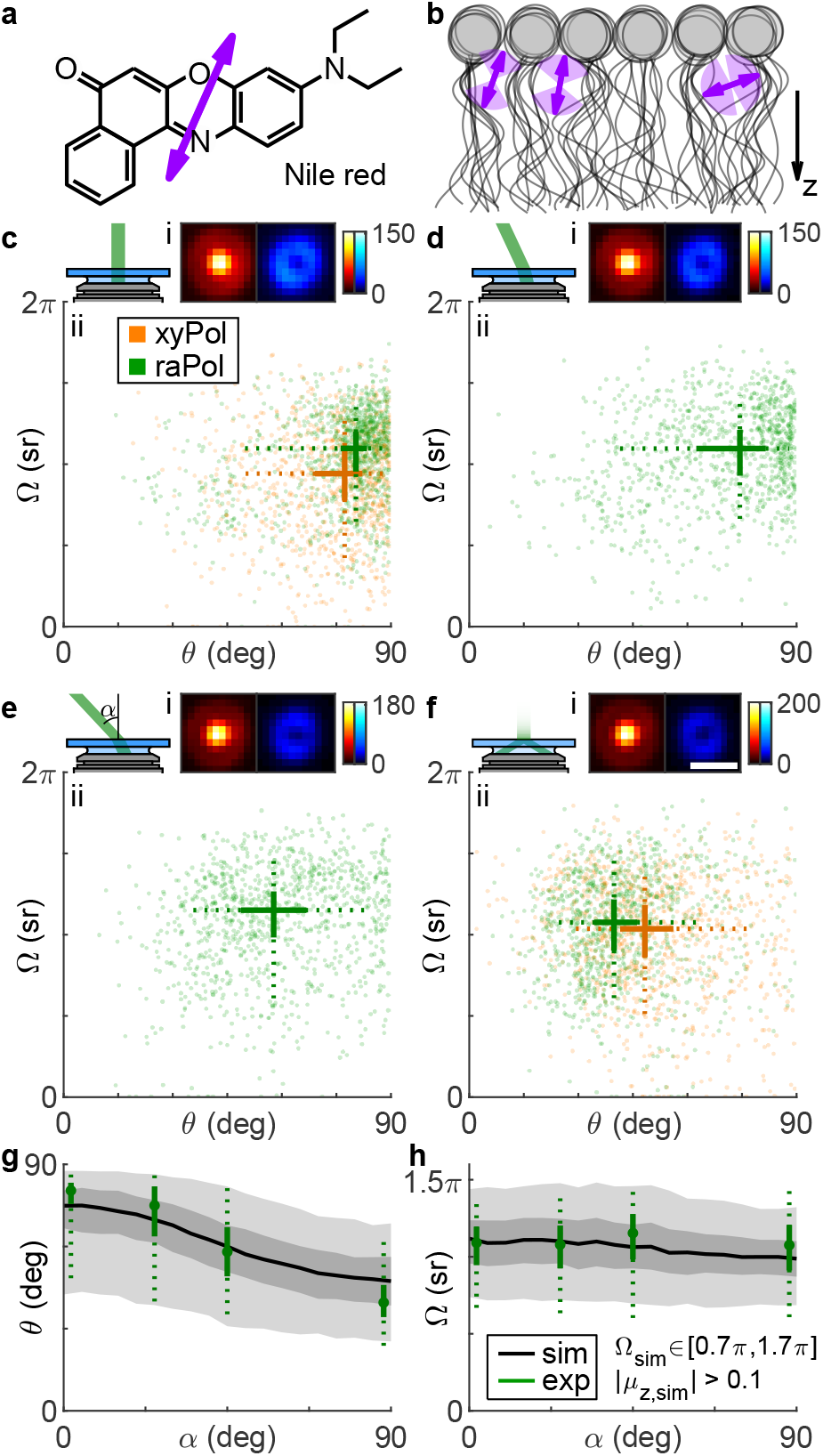
Rotational diffusion of (a) Nile red (NR) within (b) DPPC supported lipid bilayers (SLBs). Arrows represent the direction of the transition dipole moment ***μ***. ^38–40^ (c-f) (i) Average raPol images and (ii) orientation and wobble measurements of NR as a function of illumination tilt angle *α*: (c) 0°, (d) 25°, (e) 45°, and (f) total internal reflection fluorescence (TIRF). Measurements were collected using both (orange) *xy*Pol and (green) raPol imaging. Crosses represent the median of the distribution, while solid lines represent the 33rd and 67th percentiles, and dotted lines represent the 10th and 90th percentiles. (g,h) Quantifying the effect of illumination tilt angles *α* on measurements of (f) polar angle *θ* and (h) wobble Ω. Green: experimental measurements. Dots represent the median, solid lines represent 33rd and 67th percentile, and dotted lines represent the 10th and 90th percentile. Black and gray: simulated orientation measurements of rotationally diffusing SMs, assuming an excited-state lifetime of 4.6 ns, rotational diffusion coefficient of 0.035 rad^2^/ns, and constrained diffusion within a hard-edged cone of solid angle Ω_sim_ ∈ [0.7*π*, 1.7*π*] sr. Lines, dark and light gray areas represent the median, 33rd/67th, and 10th/90th percentiles, respectively. Colorbar: photons; scalebar: 500 nm.

To explain how the polarization of the illumination laser strongly affects the observed NR orientations, we used Monte Carlo simulations to model light absorp-tion, rotational diffusion, ^42,43^ and photon emission of NR within a DPPC SLB, using reported values for its excited-state lifetime (4.6 ns) and rotational diffusion coefficient (0.035 rad^2^/ns). ^44^ These simulations generate images of diffusing SMs (SI Section 4.1 and Figure S9), and we use RoSE-O to obtain measurements of their average orientation (*θ*_sim_, *ϕ*_sim_) and rotational wobble Ω_sim_ (Figure 2g,h). By simulating many possible models of rotational diffusion (Figure S10), we find that the best match to experimental measurements is produced when each NR molecule has a random average orientation ***μ***_sim_ uniformly distributed within the domain | *μ*_*z*,sim_ |*>* 0.1. In addition, each SM has a random wobble within a hard-edged cone of solid angle Ω_sim_ uniformly distributed between 0.7*π* and 1.7*π*; a molecule can freely explore all possible orientations within the aforementioned cone. With these parameters, the simulated orientation measurements match both the experimental median and variation in NR orientations reasonably well (7.8% average error in median 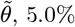 average error in median 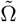).

Based on our best-fit model of rotational diffusion, we hypothesize that during a fluorescence “flash,” binding sites within the SLB constrain NR molecules such that their dipoles are oriented in all directions except parallel to the membrane (i.e., *θ* ∈[0°, 84°]). Given the angle between NR’s dipole moment and its structure (Figure 2a), there is thus a ∼54° tilt between the long axis of NR and the membrane normal. ^38–40^ Further, steric effects allow NR to explore an intermediate range of orientations (median 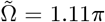 sr in our experiments irrespective of illumination tilt *α*) but prevent it from rotating freely (Figure 2b). Thus, we believe NR binds near the glycerol backbone ^44–46^ or to kinked acyl chains ^47–49^ of DPPC that are stable on the order of our camera exposure time (100 ms).

We next explore raPol’s ability to detect chol-induced ordering dynamics within DPPC SLBs. ^31^ Cholesterol is loaded into the bilayer using two successive MβCD-chol treatments (40 μM for 5 minutes, then 80 μM for 5 minutes, SI Section 3.2), and we again measured NR orientations using *xy*Pol and raPol under epi (Figure 3a) and TIRF excitation (Figure 3b). We observe two distinct “clusters” of NR orientations under both illumination conditions; the first is oriented nearly perpendicular to the SLB (median 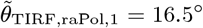), which is consistent with previous observations. ^31^ The polar angles of localizations within this cluster do not change with illumination polarization, except that number of localizations when using epi illumination is 15% of that using TIRF. Both *xy*Pol and raPol detect a decrease in rotational wobble (median 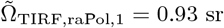 sr) when chol is present. These data suggest that this population of NR is tightly confined within a small range of orientations during its fluorescence lifetime, which is consistent with our intuition of increased crowding within cholcondensed SLBs.

**Figure 3:**
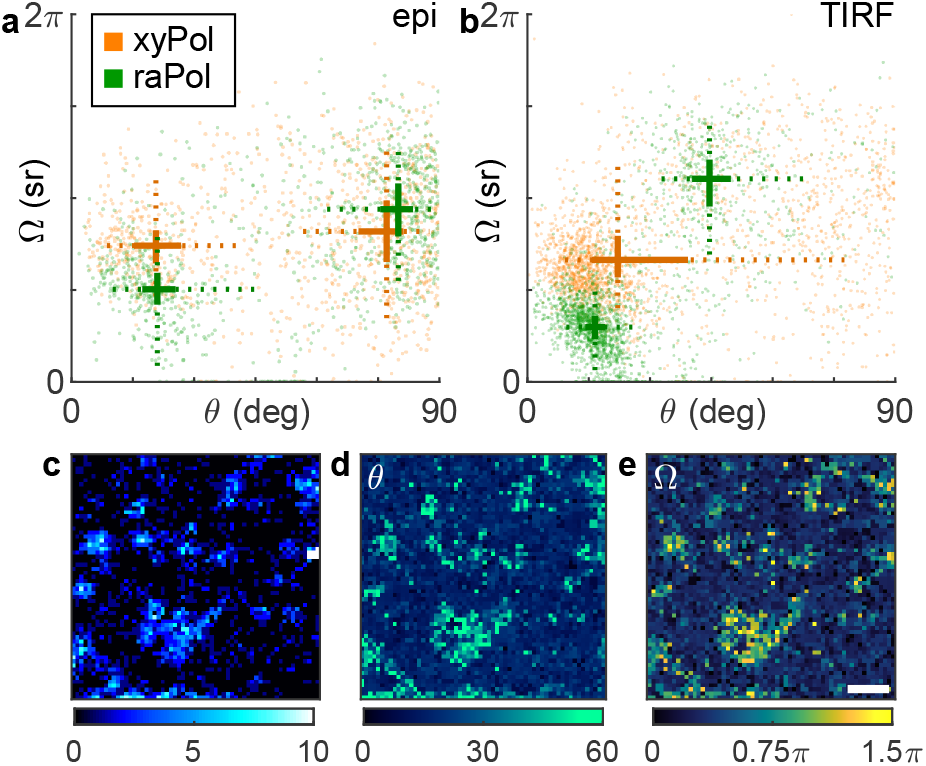
Orientation distribution of NR molecules within DPPC SLBs after cholesterol-loaded methyl-β-cyclodextrin (MβCD-chol) treatment excited by (a) epi and (b) TIRF illumination and measured using (orange) *xy*Pol and (green) raPol. Crosses represent the median of the distribution, while solid lines represent the 33rd and 67th percentiles, and dotted lines represent the 10th and 90th percentiles. A kmeans clustering (*k* = 2) is used to separate the distribution into two populations. (c-e) Identifying cholesterol-rich regions within the SLB using raPol and TIRF illumination. (c) SMLM image of NR in the azimuthally polarized imaging channel. (d-e) SMOLM images of (d) median polar angle *θ* and (e) median wobble cone angle Ω of NR localizations in each bin. Colorbars: (c) localizations per bin, (d) polar angle (deg), (e) wobble solid angle (sr). Bin size: 100 ×100 nm^2^. Scale bar: 1 μm.

However, we note a significant discrepancy between *xy*Pol and raPol measurements of the second cluster. When using raPol, this other population of NR exhibits similar orientations (median 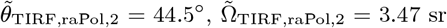 sr) to that of NR within the DPPC-only SLB (Figure 2a, 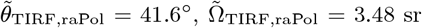 sr). However, there is no clearly resolved second cluster in the *xy*Pol measurements, just a long tail in the *θ* −Ω distribution [Figure S13a(i)]. Compared to raPol imaging (Figure 1e,f and Table S3), *xy*Pol’s degraded orientation sensitivity makes it difficult to resolve these two NR orientation behaviors within DPPC-chol SLBs.

These two distinct orientation subpopulations raise the question: are these NR binding behaviors uniformly dispersed throughout the SLB, or has chol partitioned the SLB into micro- or nanodomains? We first use ThunderSTORM to localize NR blinking events in the azimuthally polarized channel. Since NR within ordered DPPC-chol SLBs is nearly fixed and perpendicular to the membrane (Figure 3b), azimuthally polarized fluorescence only originates from regions of the SLB that are depleted of cholesterol. We observe an obvious distinction between darker cholesterol-rich regions vs. localization hotspots without cholesterol (Figure 3c). Analyzing images from both raPol channels using RoSE-O, we are able to construct SMOLM orientation *θ* (Figure 3d) and wobble Ω (Figure 3e) images, where chol-depleted nanodomains are easy to distinguish due to their large polar and wobble angles. These regions are difficult to detect in standard SMLM reconstructions since NR’s localization density is mostly uniform everywhere (Figure S13d). Interestingly, *xy*Pol is able to distinguish between chol-rich and poor regions of the SLB using NR polar angles *θ* (Figure S13b), but its map of NR wobble Ω is extremely noisy (Figure S13c).

With increasing chol in the SLB, we also observed remarkable changes in the residence time of many NR molecules. Here, we track the 2D positions and 3D orientations simultaneously of diffusing NR molecules over multiple frames (see Movie S1 for typical images of NR and Figure S16d for the distribution of NR binding times). Within the DPPC-only SLB, one molecule is translationally fixed, but rotates along the polar direction; the brightness ratio between the raPol imaging channels changes over time (Figure 4a-c, blue). After a single application of MβCD-chol, another SM exhibits more translational diffusion and rotates along the azimuthal direction; the double-spot PSF in the azimuthally polarized channel rotates dramatically over time (Figure 4a-c, orange). After the final treatment, another NR molecule moves a large distance in *x* and *y* but remains *z*-oriented throughout its entire trajectory (Figure 4a-c, purple). Additional representative SM trajectories are shown in Figure S14.

**Figure 4:**
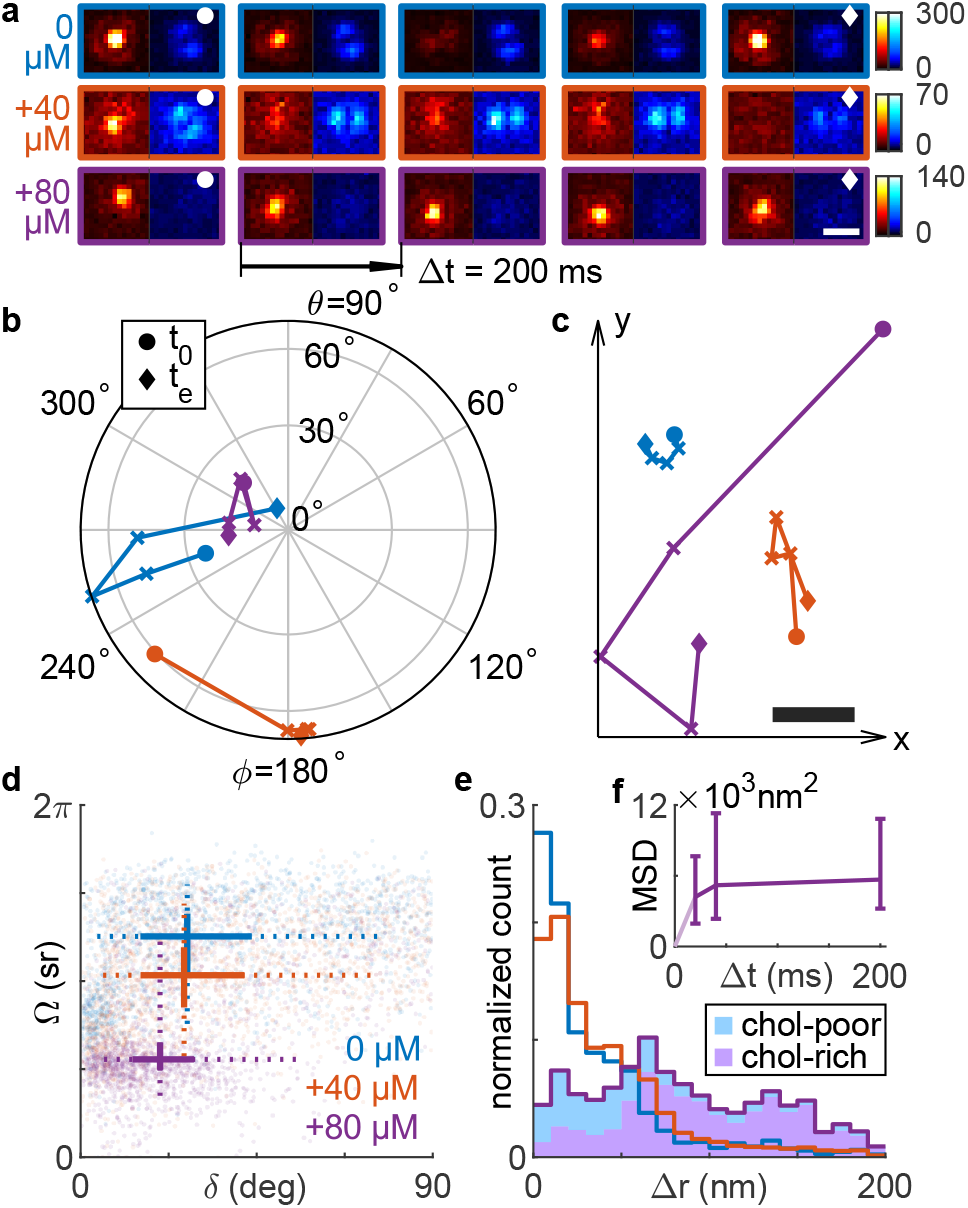
Rotational and translational dynamics of NR within SLBs before and after successive MβCD-chol treatments. Blue: before MβCD-chol treatment; orange: after 40 μM treatment for 5 minutes; purple: after 80 μM treatment for 5 minutes. (a) Raw raPol images of three SMs undergoing diffusion before and after each treatment. Scale bar: 500 nm; colorbar: photons. (b,c) SM trajectories exhibiting (b) rotational and (c) translational diffusion corresponding to images in (a). Circles and diamonds represent the first (*t*_0_) and last frame (*t*_*e*_), respectively. The relative position between the three trajectories in (c) is arbitrary. Scale bar: 50 nm. (d) Average angular displacement 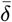 vs. av-erage wobble Ω for each NR trajectory. Solid lines represent the 33rd and 67th percentiles, and dotted lines represent the 10th and 90th percentiles of each distribution. Intersections represent the median. (e) Measured lateral displacements Δ*r* accumulated over all NR trajectories. Shaded area (purple) represents localizations within chol-rich regions of the SLB, as measured by the NR polar angle *θ* and wobble Ω. (f) Mean squared displacement (MSD, nm^2^) as a function of time lag Δ*t*. Errorbars represent the 33rd/67th percentile. All data here are from blinking events lasting at least two consecutive frames [200 ms for (a-e)].

To quantify rotational diffusion between frames, we compute the angular displacement (i.e., rotation) of single NR as

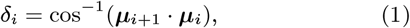

which represents an angle between two adjacent orientation measurements ***μ***_*i*_ and ***μ***_*i*+1_ within an SM trajectory. This slow-scale displacement remains unchanged after a first application of MβCD-chol [27.3° ± 26.5° (median ± std.dev.) before and 26.4° ± 25.7° after treatment, respectively, Figure 4d]. Only after a second higher-dose incubation of MβCD-chol does the average SM rotation 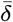 decrease to 20.3° 20.2°. In contrast, fast-scale intraframe SM wobbling Ω decreases successively after each treatment (Figure 4d), from 3.93 ±0.96 sr before treatment to 3.24 ±1.05 sr after a 40 μM incubation for 5 min., and finally to 1.74 ±1.06 sr after a 80 μM treatment.

Even though NR molecules become more rotationally constrained as chol concentration increases, they curiously exhibit successively larger translational motions. The lateral displacement, given by

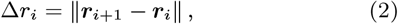

is 20.8 ± 37.4 nm (median ± std. dev., 200 ms time lag between the *i*^th^ and (*i* + 1)^th^ frame) averaged across all NR localizations before adding MβCD-chol (Figure 4e). After a 40 μM treatment, the average displacement becomes 28.2 ±37.5 nm and grows to 75.5 ±50.3 nm after a 80 μM treatment. We gain additional insight into these behaviors by using the polar orientation *θ* and wobble Ω of each localization to separate molecules exploring chol-rich regions of the SLB from those exploring chol-poor ones (using *k*-means classification, see SI Section 4.3). We find that the largest lateral displacements overwhelmingly occur when NR is within chol-rich domains [Δ*r* = 44.4 ±44.1 nm in chol-poor vs. Δ*r* = 91.5 ±47.1 nm in chol-rich domains, Figure S15c(iii)]. In fact, the proportion of all large lateral displacements (Δ*r >* 62.4 nm = 3 ×the median of Δ*r* in DPPC SLBs) that originate from chol-rich regions is more than 84.8%. Overall, however, the mean squared displacement (MSD, SI Section 4.2) of all NR trajectories measured after the 80 μM treatment scales sublinearly with the time lag Δ*t* (Figure 4f); the MSDs are (4.2 ± 13.2) ×10^3^ nm^2^, (5.2 ± 17.0) × 10^3^ nm^2^, and (5.7 ± 9.6) × 10^3^ nm^2^ for Δ*t* = 20 ms, 40 ms and 200 ms, respectively. Thus, NR diffusion is inhomogeneous and largely confined, aside from infrequent jumps, throughout the SLB.

In summary, we design and implement a new PSF, the radially and azimuthally polarized (raPol) standard PSF, for SMOLM that only requires a commercially available waveplate and polarizing beamplitter to be added to a standard epifluorescence microscope. The PSF is significantly more compact than other engineered PSFs. Despite its compactness, raPol’s orientation precision is 30.6% better on average compared to *xy*Pol. In addition, since *z*-oriented dipoles produce radially polarized light, raPol can directly detect the presence of out-of-plane molecules via a simple quantification of the brightness ratio between its two detection channels (Figure 3c). Thus, it achieves excellent sensitivity for measuring the orientation of a fluorescent molecule. ^25,26^

This combination of superior detection rate, excellent orientation measurement precision, and strong localization precision enables raPol to measure the rotational and translational dynamics of SMs with exquisite detail. In particular, raPol is able to distinguish between depolarization, i.e., fluorescence polarization that is not parallel to the excitation polarization, caused by an immobile tilted SM from an SM exhibiting large rotational diffusion–cases that are difficult to distinguish using time-resolved fluorescence anisotropy. ^50,51^ In addition, tracking the position and orientation of many SMs simultaneously within a field of view allows raPol to map dynamics within the membrane more expediently than point-scanning techniques like stimulated emission depletion (STED) FCS ^52,53^ and polarized FCS, ^54^. while also offering higher spatial resolution than ensemble measurements. ^55^ Further, since detection rate and estima-tion performance are critically important for quantitative imaging, raPol overcomes limitations of orientation-sensing PSFs, e.g., the Tri-spot and Duo-spot PSFs, ^31^ which require more photons and longer exposure times to achieve good detection performance; combining raPol with varying illumination polarizations measures NR dynamics with superior accuracy.

Our tracking of NR orientation and wobble within the membrane suggests that it binds near the glycerol backbone or to the kinked lipid tails of DPPC; due to the relative angle between NR’s transition dipole and its long axis (Figure 2a), these binding pockets are unlikely to be normal to the membrane. One possible way to corroborate our observations is to directly label the acyl chains of DPPC at various positions with bifunctional dyes ^56,57^ and compare their orientational dynamics to that of NR; while outside the scope of this work, these correlative studies would provide powerful insights into lipid conformational dynamics. ^47–49^ Moreover, the binding pockets can likely be compressed by condensing the acyl chains using cholesterol. ^31,58^ After a first dose of MβCD-chol, the average cholesterol concentration within the membrane is still relatively low (Figure S12b), but measuring the decreasing wobble Ω reveals the changing chemical composition of the SLB (Figure 4d).

After a second treatment, we find that NR demonstrates a curious combination of diffusive behaviors: strongly confined rotational motions but dramatically larger translational jumps between frames. The distribution of lateral displacements Δ*r* (Figure 4e, purple) is non-exponential and broad. NR also largely favors jumping within and between domains of similar chemical composition. We observed 16 times more jumping events between chol-rich regions versus from rich to poor; similarly, once within a chol-poor domain, NR jumped between chol-poor regions at 7.4 times the rate of jumping to chol-rich domains (SI Section 4.3). We surmise that the MβCD-chol treatments dissolve the SLB in a highly nonuniform manner, causing NR to exhibit “jump diffusion” between regions of intact membrane. NR molecules repeatedly diffuse across damaged portions of the bilayer with widely varying jump distances Δ*r* to bind transiently with other chol-rich islands, where their orientations are highly confined upon binding. Thus, NR’s rotational and translational dynamics, as observed by raPol SMOLM, support previous observations of cyclodextrin molecules damaging lipid membranes by carrying away lipid molecules after depositing cholesterol. ^59,60^ This insight into MβCD-chol activity on the SLB is not detectable by conventional SMLM (Figure S13d,e).

Looking ahead, we anticipate that the ease of implementing raPol will enable SMOLM to be adopted more widely by the nanoscience community. For example, raPol can be adapted to map nanodomain partitioning, the saturation of acyl chains, and temperature variations within SLBs; while these phenomena have been studied by tracking SM translational dynamics, ^61–63^ we hypothesize that simultaneous measurements of translational and rotational diffusion will enable more sensitive experiments when these effects are more subtle. Such studies would be aided by improvements in image analysis algorithms to increase their automation, robustness, and speed. We look forward to continued developments in SM orientation-localization microscopy for elucidating a more complete picture of chemical dynamics at the nanoscale.

## Acknowledgement

This work is supported by the National Science Foundation under grant number ECCS-1653777 and by the National Institute of General Medical Sciences of the National Institutes of Health under grant number R35GM124858.

## Supporting Information Available

Imaging system schematic and alignment and calibration procedures; characterization of raPol’s detection and estimation performance; additional NR orientation-localization analysis; additional references. ^64,65^ The data underlying this study are openly available in OSF at https://osf.io/64bfv/?view_only=3a2cbdc3fd20467486f3574d54ecc7f6 and from the corresponding author upon reasonable request.

## TOC Graphic

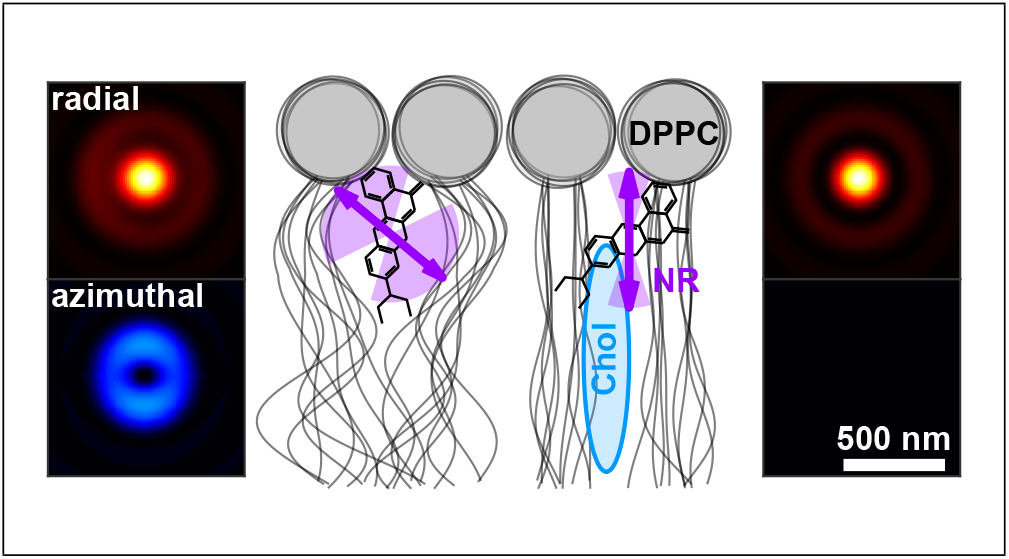

